# Pharmacological but not physiological GDF15 suppresses feeding and the motivation to exercise

**DOI:** 10.1101/2020.10.23.352864

**Authors:** Anders B. Klein, Trine S. Nicolaisen, Niels Ørtenblad, Kasper D. Gejl, Rasmus Jensen, Andreas M. Fritzen, Emil L. Larsen, Kristian Karstoft, Henrik E. Poulsen, Thomas Morville, Ronni E. Sahl, Jørn W. Helge, Jens Lund, Sarah Falk, Mark Lyngbæk, Helga Ellingsgaard, Bente K. Pedersen, Wei Lu, Brian Finan, Sebastian B. Jørgensen, Randy J. Seeley, Maximilian Kleinert, Bente Kiens, Erik A. Richter, Christoffer Clemmensen

## Abstract

Growing evidence supports that pharmacological application of growth differentiation factor 15 (GDF15) suppresses appetite but also promotes sickness-like behaviors in rodents via GDNF family receptor α-like (GFRAL)-dependent mechanisms^1,2^. Conversely, the endogenous regulation and secretion of GDF15 and its physiological effects on energy homeostasis and behavior remain elusive. Here we show, in four independent studies that prolonged, moderate- to high-intensity endurance exercise substantially increases circulating GDF15, in a time-dependent and reversible fashion, to peak levels otherwise only observed in pathophysiological conditions. This exercise-induced increase can be recapitulated in mice following forced treadmill running and is accompanied by increased *Gdf15* expression in the liver, skeletal muscle, and heart muscle. Compared to other metabolic stressors, like fasting, acute high-fat diet feeding, severe caloric excess and temperature changes, exercise has a greater impact on circulating GDF15 levels. However, whereas pharmacological GDF15 inhibits appetite and suppresses wheel running activity via GFRAL, in response to exercise, the physiological induction of GDF15 does not. In summary, exercise-induced circulating GDF15 correlates with the duration of endurance exercise. However, higher GDF15 levels after exercise are not sufficient to evoke canonical pharmacological GDF15 effects on appetite or responsible for exercise aversion/fatigue. Thus, the physiological effects of GDF15 as an exerkine remain elusive.

## Main text

GDF15 was originally discovered as a macrophage inhibitory cytokine (MIC-1)^3^ but in recent years has attracted considerable attention due to its anti-obesity potential^4^. Administration of recombinant GDF15 to rodents and monkeys promotes substantial anorexia and weight loss^5–9^. However, GDF15 administration also promotes aversive behavior and nausea in rodents^2,10^. Whereas the pharmacological effects of GDF15 have undergone considerable scientific scrutiny, the normophysiological role of endogenous GDF15 on energy homeostasis and behavior remains unknown. We have previously shown that one hour of vigorous cycling exercise increases circulating GDF15 in humans^11^. Given the role of exogenous GDF15 on malaise-related anorexia, we wondered if endogenously-derived GDF15 promotes a similar behavior. Herein, we pursue the hypothesis that exercise-induced GDF15 is an endocrine feedforward signal that protects the organism from excessive physical and/or metabolic stress by signaling exercise aversion through GFRAL. Further, we investigate the possibility that GDF15 is responsible for the temporary anorexia after intense exercise.

If GDF15 is a feedforward signal to protect the body from excessive exercise stress, then it should be secreted in response to any type of exercise of sufficient intensity and to a greater extent during prolonged exercise. To investigate this rationale, we first compared the effects of endurance vs resistance exercise on circulating GDF15 in a randomized crossover study of 1 hour of bicycling at 70% of VO2_peak_ or 1 hour of high-volume resistance exercise in moderately trained males. Both endurance and resistance exercise increased circulating GDF15 measured immediately after exercise by ~15% (**Fig. 1a**). During the 3 hr post-exercise recovery period, GDF15 plasma levels increased faster following the cycling exercise bout, but levels were similarly increased by ~25% 3 hr after both modes of exercise, suggesting that exercise increases GDF15 independent of exercise mode. We next tested circulating GDF15 concentrations in elite male triathletes cycling for 4 hr at 73% of their maximum heart rate and found a remarkable 5.3-fold increase in plasma GDF15 compared to baseline (**Fig. 1b**). More than 30% of the subjects reached plasma levels higher than 2.5 ng/ml, levels usually only seen in pathological conditions^12,13^. Of note, GDF15 levels returned to baseline values within 24 hr after exercise. In well-trained subjects, we observed a comparable 4.5-fold increase in circulating GDF15 following the completion of a marathon with an average time of 4 hr (**Fig. 1c**). To evaluate the role of exercise duration and substrate availability on exercise-induced GDF15 levels, we analyzed plasma levels of GDF15 in another group of well-trained subjects that were instructed to cycle to exhaustion under three different dietary regimes that subjects adhered to for 72 hr prior to the exercise bout: a standard diet, a low-carbohydrate diet, and a high-carbohydrate diet. A carbohydrate-rich diet, which increases muscle glycogen content, delays fatigue substantially during exercise time trials^14^. In this study, the induction of circulating GDF15 was greatest during the high-carbohydrate trial, which also coincided with the longest trial time (149 min, 4.5 fold increase in GDF15) (**Fig. 1d**). Whereas these data do not allow for making firm conclusions about whether exercise duration *per se* or certain substrate availability in combination with exhaustive exercise plays a role for the exercise-induced GDF15, they do underscore that exercise stress promotes a steep increase in GDF15 secretion once the activity goes beyond 2 hr (**Fig 1b, d, e**). Conversely, GDF15 was least induced in the low carbohydrate trial in which subjects exercised a considerably shorter time (69 min, 1.8 fold increase in GDF15). Interestingly, when subjects reached exhaustion, circulating GDF15 levels were ~1 ng/ml, considerably less than compared to ~2 ng/ml at exhaustion following high carbohydrate feeding. Together, these data underline that exercise duration is important for exercise-induced GDF15 levels. This raises the possibility that during low carbohydrate availability, comprising low muscle and liver glycogen stores, hindbrain sensitivity to GDF15 is increased in order to avoid behaviors that compromise glucose supply to the CNS. Such a central “set point” for the perceived exhaustion could involve an uncharacterized liver-hindbrain crosstalk with GDF15 as a key humoral signal.

**Figure 1.**
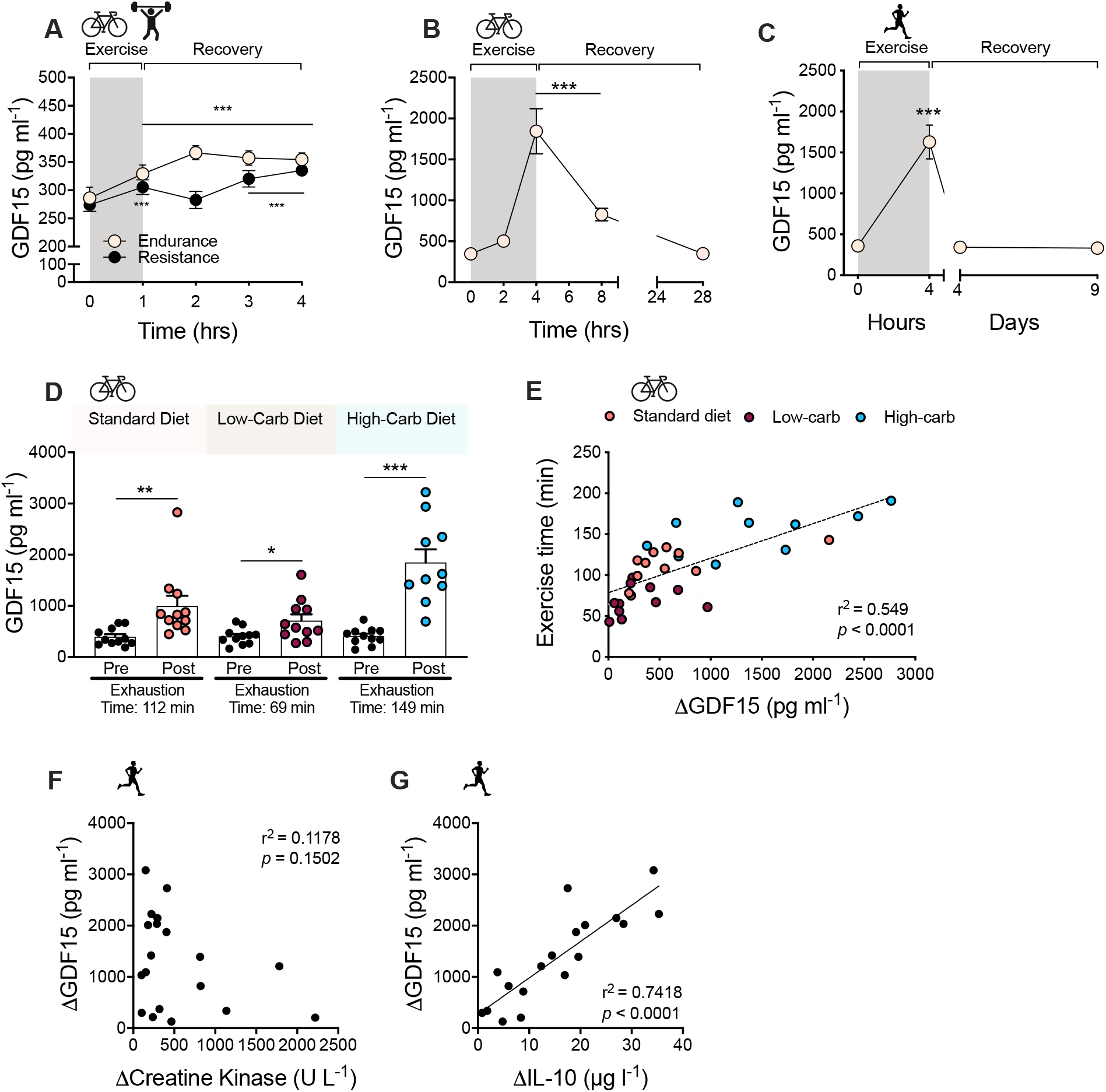
Effect of exercise on circulating GDF15 in humans. **a**, Plasma GDF15 values before and after 1 hr of high-intensity resistance (n=10) or vigorous endurance exercise (n=10) and at indicated timepoints during recovery. **b**, Plasma GDF15 leves were measured in 15 highly trained elite male triathletes bicycling at 73 ± 1% of HR_max_ for 4 hr, with blood sampling as indicated immediately before, during, and in recovery from exercise. **c**, Plasma GDF15 levels measured before and after a marathon (42,195 km), completing time was 243 +/− 8 minutes [mean ± SEM] (4 hours on the figure), with additional measurements of GDF15 on day 2-5 and day 9-10 after the marathon (on day 4 and day 9) (n = 20). **d**, On three separate occasions, under three different dietary regimes as indicated, subjects performed cycling exercise at 75% of VO_2max_ until exhaustion. GDF15 plasma levels were measured pre exercise and at exhaustion on each occasion (n=11, one missing value in the high-carb group post-exercise). **e**, Relationship between change in GDF15 values (ΔGDF15) and time to exhaustion for the diet-exercise interventions presented in **d**. **f**, Relationship between change in GDF15 values (ΔGDF15) (from **c.**) and Δcreatine kinase pre vs. post marathon. **g**, Relationship between change in GDF15 values (ΔGDF15) (from **c**) with ΔIL-10 plasma values pre vs. post marathon. Two outliers were detected and removed from **g** using automatic outlier detection (ROUT method). Data are presented as mean +/− SEM,*P<0.05; **P<0.01; ***P<0.001 compared to time-point 0 or as indicated. **a**, Repeated measures two-way ANOVA with Bonferroni multiple comparisons test. **b** and **c**, One-way ANOVA repeated measures relative to starting time, **d**, Paired t-tests for each exercise-diet trial.

A greater increase in circulating GDF15 after strenuous exercise (2 hr or more) also could be related to muscle damage. However, the prolonged cycling was performed by highly trained athletes accustomed to this type of exercise. Further, we observed no correlation between changes in levels of plasma creatine kinase, a marker of muscle damage, and changes in levels of GDF15 in response to a marathon run (**Fig. 1f**), ruling out this possibility. In a previous study, we measured GDF15 in arterial and venous blood sampled from the exercising leg and found similar GDF15 concentrations in both vessels, suggesting that GDF15 is not released from skeletal muscle during exercise in humans^11^. Therefore, other organs likely secrete GDF15 during exercise. IL-6 is a well-established myokine with pleiotropic whole body effects that is rapidly released from skeletal muscle during exercise^15^. To probe if exercise-induced IL-6 secretion from muscle could trigger GDF15 release in other organs, we investigated plasma GDF15 in healthy humans infused with two doses of recombinant human IL-6 followed by a mixed meal test (**Extended Data Fig. 1a**). Administration of IL-6 alone or in combination with a mixed meal did not have impact on circulating levels of GDF15. Another major anti-inflammatory cytokine induced by exercise is IL-10^16^. The induction of IL-10 in response to exercise is delayed compared to that of IL-6, and the magnitude of exercise-induced IL-10, like GDF15, is related to the duration of exercise^17^. Notably, we observed a strong correlation between the exercise-mediated increase in circulating IL-10 and exercise-induced GDF15 following a marathon run (**Fig. 1g**). Whether there is a causal relationship between exercise-induced GDF15 and IL-10 remains to be determined. In summary, these data, from four independent human exercise studies, clearly underscore that different types of exercise increase circulating GDF15 levels and, with longer bouts of moderate-to high intensity exercise, GDF 15 levels increase 4-5-fold.

It was recently demonstrated that an increase in circulating GDF15, in response to metformin treatment, promotes weight loss in a GFRAL-dependent fashion^18,19^. Given that prolonged exercise leads to circulating GDF15 levels comparable to the GDF15 levels reported following metformin treatment, it raises the intriguing possibility that exercise-induced increases in endogenous GDF15 mediate the effects of intense exercise on transient food intake suppression^20,21^. To address this possibility mechanistically in mice, we established a forced running protocol on an inclined treadmill that resulted in a >4 fold increase in plasma GDF15 levels post exercise (**Fig. 2a**), similar to what we observed with longer duration, high intensity exercise in humans (**Fig. 2b**, **Fig. 1c**). Immediately after this treadmill exercise paradigm, *Gdf15* mRNA was increased 2-fold in liver, heart and soleus muscle, suggesting that key tissues involved in substrate metabolism during exercise are contributing to the increased circulating GDF15 levels (**Fig. 2c,d,e**). The increased expression of *Gdf15* in these tissues coincided with an increase in markers related to the unfolded protein response (UPR) cellular stress pathway such as *Xbp1*, *Atf4*, and *Atf6* (**Fig. 2c,d,e**). Important to note, basal *Gdf15* expression is substantially higher in the liver compared to skeletal muscle and heart muscle (24-fold and 14-fold, respectively) (**Extended data Fig. 2a**). Thus, despite that an increase in *Gdf15* mRNA in human exercising muscle has been reported^22^, it remains unclear if GDF15 is a relevant myokine^11^. In contrast to forced treadmill running, voluntary wheel running does not increase circulating GDF15 levels, likely due its lower intensity and/or intermittent nature (**Fig. 2f**).

**Figure 2.**
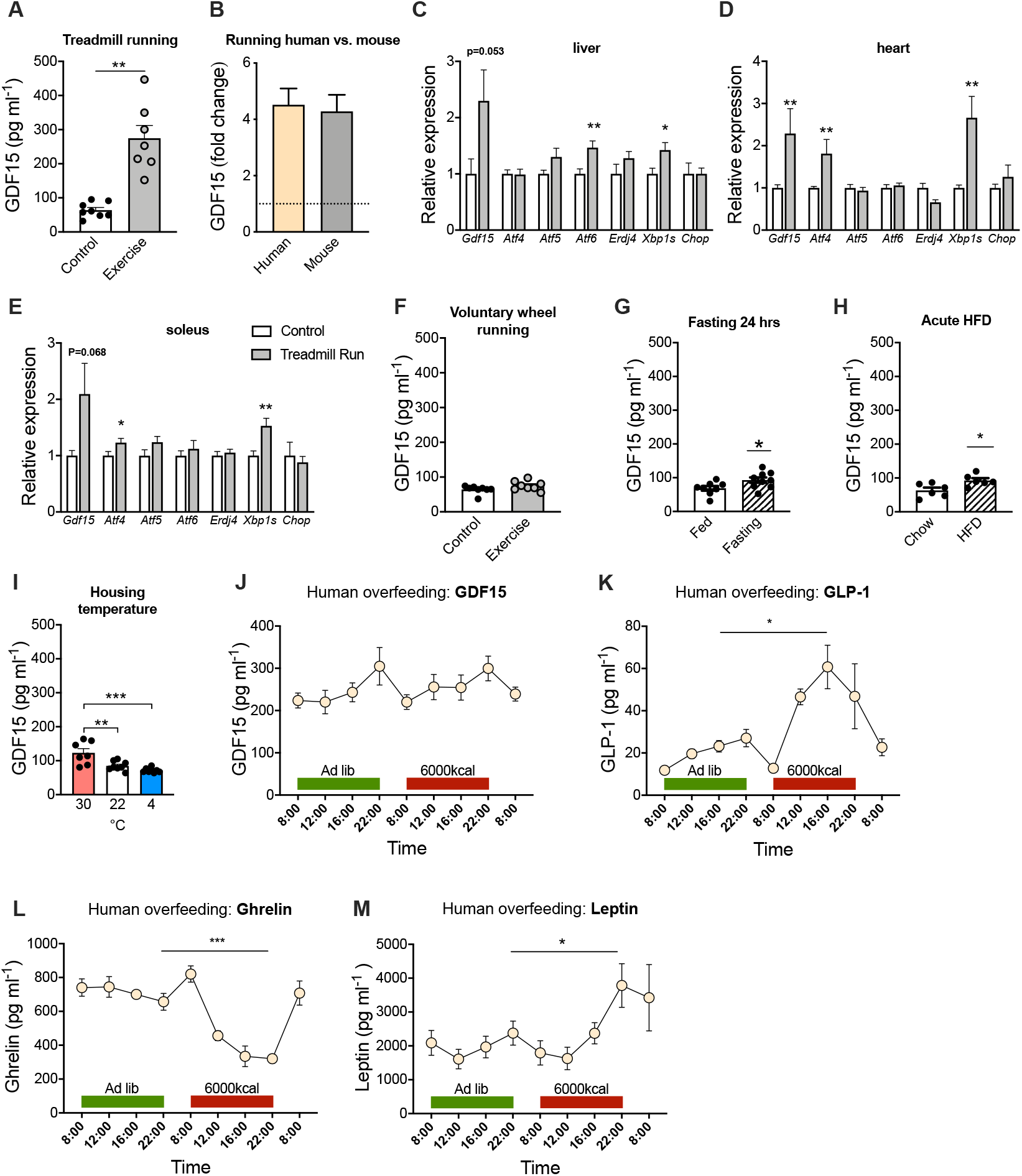
Effect of physiological metabolic stressors on circulating GDF15 in mice and humans. **a**, Plasma GDF15 levels were measured in mice exposed to forced treadmill running until exhaustion (n=8) compared to cage-matched sedentary controls (n=8). **b**, Fold-change in circulating GDF15 after exhaustive endurance running in mice (based on Fig. 2a) versus humans (based on Fig. 1c). **c, d, and e**, Relative mRNA levels of indicated genes in liver, heart muscle, and skeletal muscle (soleus) after forced treadmill running until exhaustion (n=8) compared to cage-matched sedentary controls (n=8). **f**, Circulating GDF15 levels in mice following 3 weeks of voluntary running vs. non-running controls (n=8) measured in plasma sampled 2 hr into the dark phase **g**, Circulating GDF15 levels in mice after 24 hr fasting compared to ad libitum fed mice (n=8-9). **h**, Circulating GDF15 levels in mice in response to 24 hr of ad libitum, high-fat diet exposure compared to aged-matched control mice kept on chow-fed (n=6). **i**, Circulating GDF15 levels in mice acclimatized to thermoneutrality (30°C), kept at room temperature (22°C), or mice exposed to chronic cold (4°C) for 3 weeks (n=8). **j**, **k**, **l**, **m**, Plasma GDF15, GLP-1, ghrelin, and leptin concentrations at indicated time points before, during, and after short-term overfeeding (6000 kcal in 14 hr) in young healthy male human subjects (n=5). Data are presented as mean +/− SEM,*P<0.05; **P<0.01; ***P<0.001. **a-h**, Unpaired t-test. **i**, One-way ANOVA repeated measures. **j-m**, Unpaired t-test between peak time-point for OF vs. corresponding ad lib time-point.

A high intensity exercise bout represents a transient physiological and metabolic stressor. To understand whether other forms of physiological and metabolic stressors trigger similar increases in circulating GDF15 levels, we assessed the effects of fasting, acute high-fat diet (HFD) feeding, and changes in ambient temperature on circulating GDF15 in mice and then evaluated the effect of acute, severe overfeeding in human subjects. Fasting for 24 hr increased plasma GDF15 levels by 35% in mice (**Fig. 2g**). A change from chow diet to HFD resulted in 48% increase in plasma GDF15 levels 24 hr following the diet switch (**Fig. 2h**). Mice acclimatized to thermoneutral conditions (30°C) had 45% higher GDF15 levels compared to mice kept at standard housing temperature (22°C), whereas chronic cold stress (4°C), relative to 22°C, had no effect on circulating GDF15 levels (**Fig. 2i**).

As a stress-responsive hormone, GDF15 might protect against severe overfeeding (OF), a nutritional stressor, via signaling food aversion^10^. To test this theory, we measured plasma GDF15 levels in healthy humans before, during, and after they consumed 6000kcal within 14 hr. GDF15 levels showed a diurnal pattern with peak values around 10 pm, which is consistent with findings by Tsai et al^23^. However, severe short-term OF failed to elevate circulating GDF15 (**Fig. 2j**), arguing against a role of GDF15 in short-term physiological energy balance regulation. In contrast, canonical appetite hormones, GLP-1 (160% increase at 4pm OF vs. 4pm *adlibitum*), ghrelin (50% decrease at 10pm OF vs. 10pm *ad libitum*), and leptin (60% increase at 4pm OF vs. 4pm *ad libitum*) were markedly affected by OF (**Fig. 2k, l, m**). Overall, these canonical stressors of energy homeostasis affected circulating GDF15 levels to a much lesser extent than exercise in mice and humans, suggesting that vigorous exercise is especially effective for promoting acute GDF15 secretion.

To investigate if GDF15 modulates subsequent exercise performance or exercise behavior, we tested a single pharmacological dose of recombinant human GDF15 (rhGDF15) in mice (**Fig. 3a**) exposed to either controlled treadmill running or voluntary wheel running. Administration of rhGDF15, at a dose that suppressed acute feeding by ~50% in obese mice (**Extended Data Fig. 3a**), had no influence on treadmill running to exhaustion in mice (**Fig. 3b**), suggesting that rhGDF15 has no adverse effects on the neuromuscular system or on cardiovascular fitness *per se*. In contrast, administration of rhGDF15 substantially lowered voluntary wheel running in mice adapted to running wheels (**Fig. 3c**). Decreases in voluntary running by rhGDF15 administration was sustained over a once daily treatment study for 7 days, by an average of −1.4km/day (**Fig. 3d**). Notably, these effects on voluntary running were dissociated from the anorectic effects of rhGDF15 administration, which only induced a transient decrease in food intake with no effect on body weight (**Fig. 3e**) (**Extended Data Fig. 3b)**. This implies that the effect of rhGDF15 on voluntary running behavior is more potent than the anorexigenic effects of rhGDF15. Our observation agrees with findings of a previous study showing that GDF15 administration more potently induces taste aversion compared to appetite suppression^10^. The exercise aversion following rhGDF15 treatment was absent in mice lacking the GDF15 receptor, GFRAL (**Fig. 3f, g**), underscoring that this behavior is mediated via the canonical GDF15-GFRAL hindbrain signaling pathway. Together these data emphasize that pharmacological GDF15 potently suppresses voluntary exercise, and this is unrelated to an effect on treadmill exercise capacity. These findings extend the reported effects of pharmacological GDF15 on sickness-like behaviors, including taste aversion^10^, nausea, and emesis^2^.

**Figure 3.**
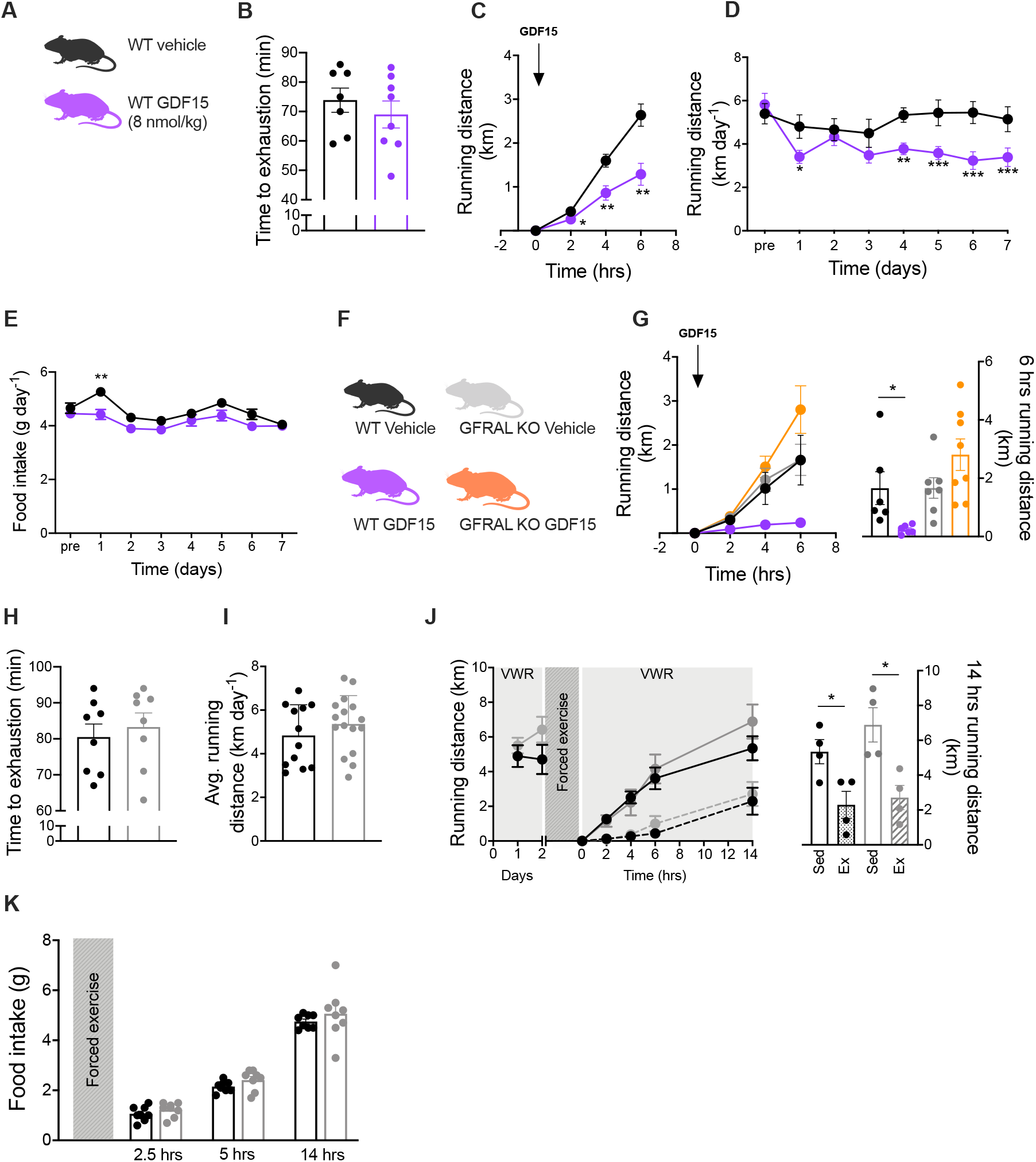
Effects of pharmacological GDF15 and exercise-induced GDF15 on food intake and running behavior. **a, b**, Effect of single subcutaneous injection of rhGDF15 (8 nmol/kg, n=8) or vehicle (n=7) on forced treadmill running until exhaustion in mice. **c**, Effect of single subcutaneous injection of rhGDF15 (8 nmol/kg, n=8) or vehicle (n=8) administered at the onset of the dark cycle on voluntary wheel running distance. **d, e** Effect of daily subcutaneous injections (at the onset of the dark cycle) of rhGDF15 (8 nmol/kg bw, n=16) or vehicle (n=16) on voluntary wheel running (d) and food intake (e). **f**, **g**, Effect of single subcutaneous injection of rhGDF15 (8 nmol/kg) or vehicle on voluntary wheel running in WT (vehicle n=6; rhGDF15 n = 6) and GFRAL KO (vehicle n=7; rhGDF15 n=8) mice. **h**, Forced treadmill running to exhaustion in WT (n=8) and GFRAL KO (n=8) mice. **i**, Voluntary wheel running distance in WT (n=12) and GFRAL KO (n=16) mice. **j**, Effect of forced exhaustive treadmill exercise on voluntary wheel running (VWR) in WT (n=4 sedentary vs. n=4 forced exercise) GFRAL KO (n=4 sedentary vs. n=4 forced exercise) mice. **k**, Effect of forced exhaustive treadmill exercise on food intake in WT (n=8) GFRAL KO (n=8) mice. Data are presented as mean +/− SEM,*P<0.05; **P<0.01; ***P<0.001. **b, g, h, i, j, k**, Unpaired t test. **c**, **d**, **e** Repeated measures two-way ANOVA with Bonferroni multiple comparisons test.

To test the hypothesis that an *endogenous* increase in circulating GDF15 levels by controlled strenuous exercise is a physiological stressor that triggers aversive circuits via the brainstem to discontinue the activity, we exercised WT and GFRAL KO mice using forced treadmill running, after which mice were given free access to running wheels in their home cages. Prior to administering the treadmill running bout, we observed that exercise capacity and baseline voluntary wheel running activity were similar between WT and GFRAL KO mice with both groups running ~5 km/day (**Fig. 3g, h, i**). Forced treadmill running substantially decreased voluntary running for several hours post exercise in both WT and GFRAL KO mice, indicating that this behavioral effect was independent of the endogenous GDF15-GFRAL signaling axis (**Fig. 3j**). Likewise, post exercise food intake was also similar between WT and GFRAL KO mice, indicating that forced exhaustive treadmill running failed to elicit GFRAL-dependent effects on appetite (**Fig. 3k**).

The data presented here identify GDF15 as a physiological, exercise-induced, stress signal. Given the emerging understanding that the weight-lowering effects of pharmacological GDF15 cannot be dissociated from sickness-like behaviors^2^, we experimentally pursued the notion that endogenously-induced GDF15 in response to exercise stress contributes to exercise fatigue as a strategy to protect the organism and preserve energy. In line with this, we discovered that vigorous endurance exercise, lasting for more than 2 hours, elicits circulating GDF15 levels comparable to those observed in patients suffering from mitochondrial disease^13^, infection^12^, and cancer^24^. Such pathological conditions are frequently accompanied by sickness-related anorexia and weight loss, and there is evidence linking GDF15 to non-homeostatic weight loss^2,6^. It is notable that the animal data presented here confirm that pharmacological GDF15 indeed inhibits feeding and reduces the motivation to exercise. In contrast, however, physiological induction of *endogenous* circulating GDF15 in response to strenuous exercise does not appear to be responsible for exercise-related aversion or suppression of food intake. Future studies are warranted to uncover this dissociation of the metabolic and behavioral consequences of pharmacological versus physiological GDF15 and to define the role of GDF15 as an exerkine.

## Methods

### Human studies

#### Study 1: Endurance versus resistance exercise

Ten healthy, young, moderately trained males participated in the study (age, 24±1 years; BMI 23.7±0.6 kg m^−2^; peak oxygen uptake (VO_2peak_) 55.7±1.7 ml min^−1^ kg^−1^, [mean ± SEM]). All subjects gave written consent to take part of the experimental procedures. Study methods have been described in detail previously^25^. Briefly, on two occasions separated by 1 week, subjects were instructed to cycle at 70% of VO_2peak_ for 1 hr or follow a high-volume resistance exercise program for 1 h. Venous blood samples were drawn before exercise, immediately post-exercise and at multiple time-points post exercise. The study was approved by the local ethics committee of Copenhagen and Frederiksberg (H-17030450) and conformed to the standards described in the Helsinki Declaration.

#### Study 2: Exhaustive cycling in elite endurance athletes

Fifteen elite male triathletes (27.0 ± 0.8 years; VO_2peak_ 66.1 ± 1.3 ml min^−1^ kg^−1^ [mean ± SEM] were fully informed of any risks associated with the study before providing informed verbal and written consent. All experimental procedures have been described in detail elsewhere^26^. In short, participants received a standard diet before cycling at 73±1% of their maximum heart rate for 4 hr in the laboratory. During the first 4 hr of recovery, subjects received either water or carbohydrate. Venous blood samples were drawn before exercise, after 2 hr of exercise, and immediately post-exercise. Additional blood sampling was done 4 hr and 24 hr post exercise. The study was approved by the ethics committee of the Region of Southern Denmark (Project ID: S-20090140) and conformed to the standards described in the Helsinki Declaration.

#### Study 3: Marathon run

Twenty healthy, moderately trained males were included in the study (age, 32.0 ± 8.5 years; BMI 24.7 ± 2.0 kg m^−2^; VO_2peak_ 52.5 ± 4.6 ml min^−1^ kg^−1^ [mean ± SD]). Venous blood samples were collected within the week before to the marathon and immediately after completion of the run. Additional sampling was done approximately 4 (2-5) and 9 (9-10) days after the marathon. The study was an observational study conducted in relation to Copenhagen Marathon 2018 and was approved by the Biomedical Ethical Committee of the Capital Region of Denmark (H-17041877) and conformed to the standards described in the Helsinki Declaration.

#### Study 4: Cycling to exhaustion on different dietary regimes

Eleven young, healthy males (age 25±4 years; VO_2peak_ 59.4±7.3 ml min^−1^ kg^−1^ [mean ± SD]) were included in the study. All subjects gave their written consent before entering the study. Briefly, on three occasions, each separated by 72 hr, subjects were instructed to cycle at 75% of VO_2peak_ until exhaustion. For the 72 hr leading up to the trials, subjects ingested either a standard diet, a low-carbohydrate diet (range: 0.2-0.3 g kg BM^−1^ day^−1^), or a high-carbohydrate diet (range: 7.4-9.7 g kg BM^−1^ day^−1^). All diets were isocaloric. Venous blood samples were taken pre- and post-exercise on each occasion. The study was approved by The Regional Committees on Health Research Ethics for Southern Denmark (S-20170198) and conformed to the standards described in the Helsinki Declaration.

#### Study 5: IL-6 infusion study

Seven young and healthy males (age 24 ± 1.1 years; BMI 24.1 ± 2.4 kg m^−2^ [mean ± SD]) participated in the study. All subjects gave their written consent before entering the study. Parts of this study have previously been published and more details can be found in Lang Lerskov *et al.*^27^. On three different occasions, each separated by 7 days, subjects underwent 30 min infusion of either saline (placebo), low dose (1.5μg) recombinant human (rh) IL-6, or high dose (15μg) rhIL-6 through the antecubital vein. Immediately after infusion, a 2 hr liquid mixed meal tolerance (15 E% fat, 20E% protein and 65 E% carbohydrate) test (MMTT) was performed. Fasted venous blood samples were obtained before (0) and after (30 min) the infusion and at time points 60 and 120 min relative to the start of the MMTT. The study was approved by the ethical committee of the Capital Region of Denmark and conformed to the standards described in the Helsinki Declaration.

#### Study 6: Acute severe overfeeding

Five healthy, lean males (age 33 ± 5.3 years; BMI 23.1 ± 1.1 kg m^−2^ [mean ± SD]) were recruited to the study. The purpose of the acute overfeeding study, together with potential risks and inconveniences, was explained to all participants, and all gave signed consent prior to inclusion. Subjects were followed on three consecutive days. Subjects consumed their habitual diet on day 1 and on day 2 were exposed to a 6000kcal hypercaloric diet (37 E% fat, 12 E% protein and 51 E% carbohydrate). Venous blood was sampled at 8:00am, 12:00am, 4:00pm, and 10:00pm at day 1 and day 2 and at 8.00am on day 3 after the hypercaloric diet. The study only required an informal approval from the regional research ethics committee and conformed to the standards described in the Helsinki Declaration.

### Animal studies

#### Animals

Wild-type (WT) C57BL/6J male and female mice were obtained from (Janvier, FR). The GFRAL knockout (KO) and WT littermates were generated as previously reported^28^. One week before experimental treatment, mice were acclimatized to single housing. All experiments were done at 22°C with a 12:12-hr light-dark cycle. Mice had free access to water and chow diet (Altromin 1324, Brogaarden, DK), or when indicated a high-fat diet (D12331; Research Diets). All experiments were approved by the Danish Animal Experimentation Inspectorate (2018-15-0201-01457).

#### Forced treadmill running

One week before the experimental day, 12-14 weeks old male mice were familiarized to treadmill (TSE Systems, D) three times (10 min at 0.17 m/s), and subsequently exposed to a running paradigm of 10 min at 10 m·min^−1^ then 40 min at 27 m min^−1^ (50% of max speed) with a slope at 10° followed by gradually increased speed (2 m min^−1^) until exhaustion. Exhaustion was defined when mice fell back to the grid three times within 30 s. Mice were sacrificed immediately after exercise. Blood was immediately centrifuged at 5000 x g at 5°C for 10 min and stored at −80°C for later analyses. Tissues were rapidly frozen on dry ice and stored at −80°C for later analyses.

#### Voluntary running

Eight-ten weeks old male mice were single-housed in cages equipped with running wheels (23 cm in diameter, Techniplast, I). The amount of bedding was reduced in order to avoid wheel blockade. Running distance was measured by a computer (Sigma Pure 1 Topline 2016, D), and, after one week of familiarization to the wheels, running distance was measured daily for 3 weeks. After 3 weeks, blood samples from the tail were taken two hours after onset of the dark cycle.

#### Fasting and acute HFD-feeding

Eight weeks old male mice were single housed and fed a regular chow diet. For the fasting study, mice were randomized to either fasting (24 hr) or remaining on chow diet. For the HFD-switch study, chow fed mice were randomized to either staying on chow diet (24 hr) or having free access to a HFD for 24 hr. Blood samples from the tail were taken for measuring GDF15 concentration.

#### Housing Temperature

Eight-ten weeks old male mice fed on chow diet were randomized to housing at 30°C, at 4°C, or remained at 22°C. After 3 weeks, blood from the tail was drawn for measuring GDF15 concentration.

#### In vivo pharmacology and exercise motivation

For the exercise treadmill exhaustion test, 12-14 weeks old female WT mice were subcutaneously administered recombinant human (rh) GDF15 (Novo Nordisk) at 8 nmol kg^−1^ (5 μl per gram body weight) or vehicle 2 hr before treadmill running (identical to above descripted protocol). For the effect of rhGDF15 on voluntary wheel running, after one week of familiarization to running wheels, 8-10 weeks old female WT mice were randomized to receive either subcutaneous injection of rhGDF15 (8 nmol kg^−1^) or vehicle once daily for 7 consecutive days. Injections was done each day within the last 60 min of the light cycle. After the first dosing running distance was monitored at 2hr, 4hr, and 6hr, and, for the remaining part of the study, distance, body weight and food intake were measured once daily. For assessment of GFRAL-dependent effects of rhGDF15 on voluntary wheel running, 12-14 weeks old male WT and GFRAL KO mice were subcutaneously administered rhGDF15 at 8 nmol kg^−1^ (5 μl per gram body weight) or vehicle control prior to gaining access to running wheels. Distance was measured 2hr, 4hr, and 6hr following the pharmacological intervention Exercise time to exhaustion in WT versus GFRAL KO mice was performed according to the protocol described above. Average running distance in WT and GFRAL KO was measured in 10-16 weeks old male mice with free access to running wheels over 2 weeks.

#### Forced exercise: Subsequent effects on voluntary exercise and food intake

Eight-fourteen weeks old male WT and GFRAL KO mice were exercised to exhaustion on a treadmill (as described previously) immediately before the beginning of the dark cycle after which they were returned to their cages with running wheel, for which they had previously been adapted. Running distance was monitored at 2hr, 4hr, 6hr, and 14hr post-exercise.

In a separate experiment, the same forced exercise protocol was applied, and animals were returned to their home cages with no running wheels. Assessment of food intake was done at 2.5hr, 5hr, and 14hr post-exercise.

### Biochemistry

GDF15 was measured in human plasma or serum using the Quantikine ELISA Human GDF-15 Immunoassay (ELISA, R&D systems, catalog no. DGD150). IL-10 was determined in single determinations using multi-spot immunoassay (V-PLEX, Meso Scale Discovery, Rockville, MD, USA) on EDTA treated plasma. CK was determined in lithium-heparin treated plasma using Cobas. In mice, plasma samples were analyzed using Quantikine ELISA Mouse GDF15 Immunoassay (ELISA, R&D systems, catalog no. MGD150). The ELISA assays were used according to the protocol provided by the manufacturer.

### RNA extraction & cDNA synthesis

Tissue was quickly dissected and frozen on either dry ice or liquid nitrogen and stored at −80C. Tissue was homogenized in a Trizol reagent (QIAzol Lysis Reagent, Qiagen) using a stainless steel bead (Qiagen) and a TissueLyser LT (Qiagen) for 3 min at 20 Hz. Then, 200 μl chloroform (Sigma-Aldrich) was added and tubes shaken vigorously for 15 sec and left at RT for 2 min, followed by centrifugation at 4°C for 15 min at 12,000 x g. The aqueous phase was mixed 1:1 with 70% ethanol and further processed using RNeasy Lipid following the instructions provided by the manufacturer. For muscle tissue, the lysis procedure described the enclosed protocol in the Fibrous Tissue Mini Kit (Qiagen) was followed. After RNA extraction, RNA content was measured using a NanoDrop 2000 (Thermo Fisher) and 500 ng of RNA was converted into cDNA by mixing FS buffer and DTT (Thermo Fisher) with Random Primers (Sigma-Aldrich) and incubated for 3 min at 70°C followed by addition of dNTPs, RNase out, Superscript III (Thermo Fisher) and placed in a thermal cycler for 5 min at 25°C, 60 min at 50°C, 15 min at 70°C, and kept at −20°C until further processing.

### qPCR

SYBR green qPCR was performed using Precision plus qPCR Mastermix containing SYBR green (Primer Design, #PrecisionPLUS). Primer sequences (**Extended Data Table 1**). qPCR was performed in 384-well plates on a Light Cycler 480 Real-Time PCR machine using 2 min preincubation at 95°C followed by 45 cycles of 60 sec at 60°C and melting curves were performed by stepwise increasing the temperature from 60°C to 95°C. Quantification of mRNA expression was performed according to the delta-delta Ct method.

### Statistical analyses

Statistical analyses were performed in Graphpad Prism version 8. For comparing multiple groups, one- or two-way ANOVA with Bonferroni posthoc multiple comparisons test was used. When comparing two groups, a Students t-test was used. Unless otherwise stated, all data are presented as mean ± SEM. P < 0.05 was considered statistically significant.

## Data availability

All data that support the findings of this study are available from the corresponding author upon reasonable request.

## Acknowledgements

The study was supported by Lundbeckfonden (R164-2013-16132), the Ministry of Culture Denmark (FPK.2017-0013, TKIF2009-052), the Augustinus Foundation (17-4211), the Danish Diabetes Academy, Funded by the Novo Nordisk Foundation, Independent Research Fund Denmark, TrygFonden and Fonden til Lægevidenskabens Fremme – AP Møller Fonden. RJS received funding from the NIH (P0I DKK117821 and R01 DK119188). EAR is supported by the Novo Nordisk Foundation grants NNF17OC0027274 and NNF18OC0034072. BK is supported by research project grants from the Novo Nordisk Foundation (19OC0057404) and the Danish Council of Independent Research - Medical Sciences. CC is supported by research grants from the Lundbeck Foundation (Fellowship R238-2016-2859) and the Novo Nordisk Foundation (grant number NNF17OC0026114). Novo Nordisk Foundation Center for Basic Metabolic Research is an independent Research Center, based at the University of Copenhagen, Denmark, and partially funded by an unconditional donation from the Novo Nordisk Foundation (www.cbmr.ku.dk) (Grant number NNF18CC0034900).

## Supplemental Material

**Extended Data Fig. 1.**
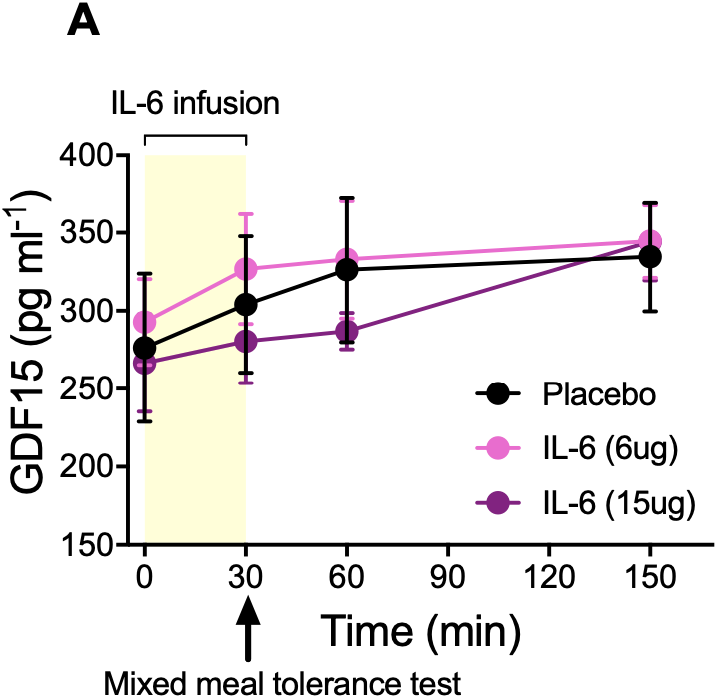
Effect of rIL-6 infusion in together with a mixed meal tolerance test on circulating GDF15 levels in healthy humans. **a**, Plasma GDF15 concentration during IL-6 infusion followed by a mixed meal tolerance test, n=7. Data are presented as means ± SEM.

**Extended data Fig. 2.**
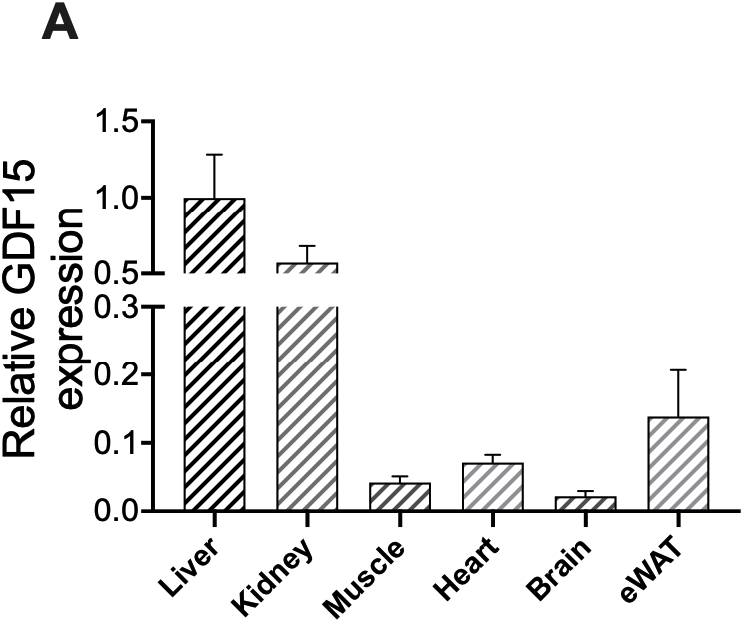
*Gdf15* expression in mice. **a**, Mouse tissue panel of *Gdf15* expression, n=8. Data are presented as means ± SEM.

**Extended data Fig. 3.**
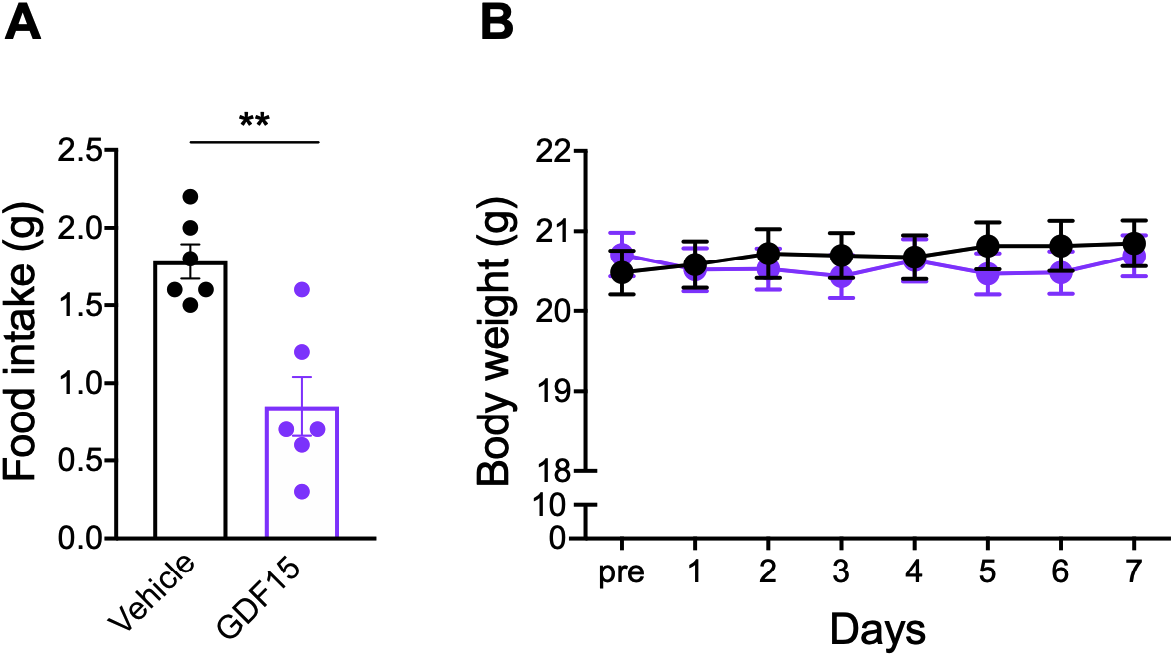
Effect of rhGDF15 administration on acute feeding in DIO mice, and during a 7 day running study in chow-fed mice. **a**, Overnight food intake of diet-induced obese (DIO) mice acutely treated with vehicle (n=6) or rhGDF15 (n=6), Body weight of lean mice treated daily with vehicle (n=6) or rhGDF15 (n=6) while having free access to running wheels. Data are presented as means ± SEM. ** P < 0.01, unpaired t-test.

**Table 1.**
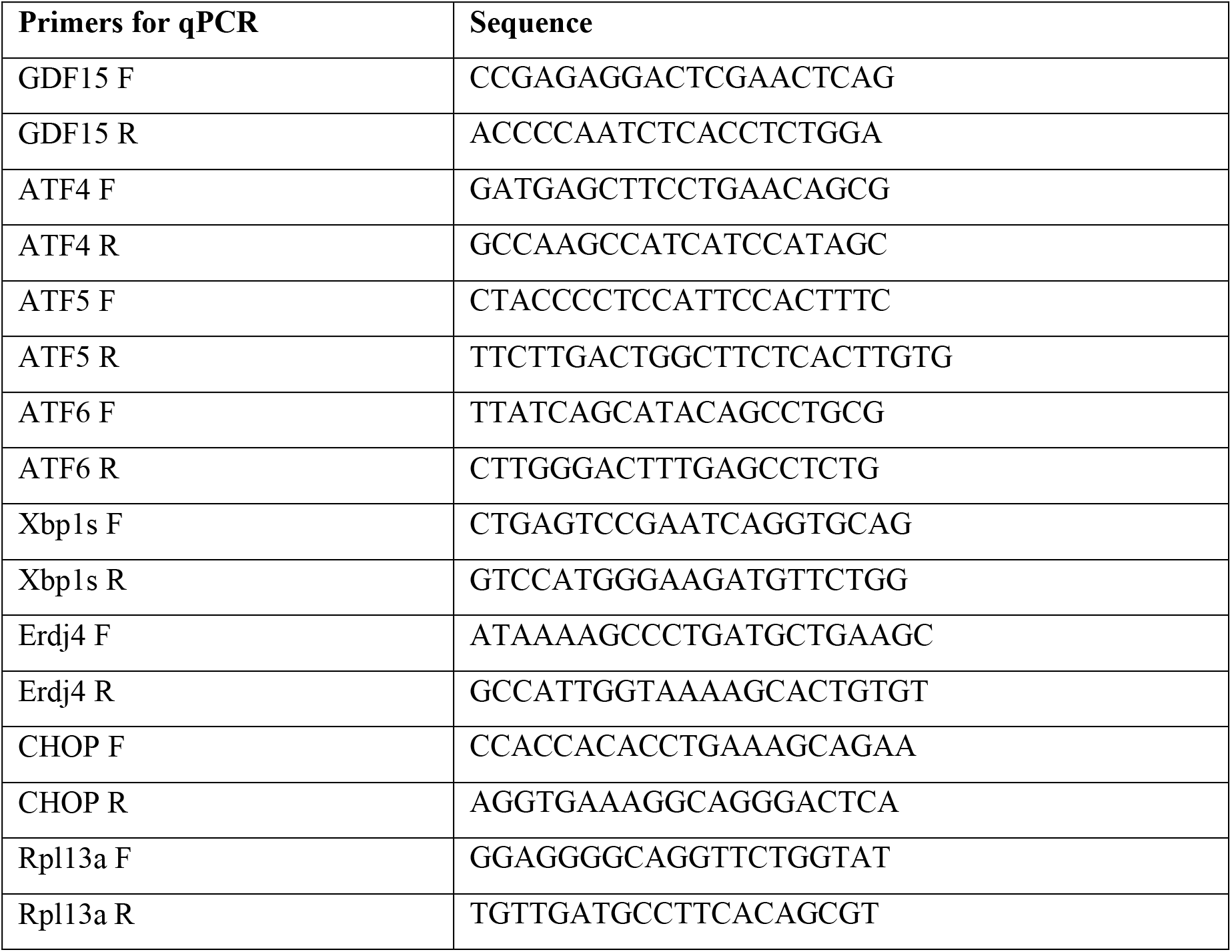
Primers for RT-qPCR.

